# Relationships of trainee dentists’ empathy and communication characteristics with simulated patients’ assessment in medical interviews

**DOI:** 10.1101/407346

**Authors:** Sho Watanabe, Toshiko Yoshida, Takayuki Kono, Hiroaki Taketa, Noriko Shiotsu, Hajime Shirai, Yasuhiro Torii

## Abstract

**Objectives:** We aimed to clarify the characteristics of communication between trainee dentists and simulated patients (SPs) and to examine how the level of trainee dentists’ self-reported empathy influences assessment by SPs in medical interviews.

**Materials and methods:** The study involved 100 trainee dentists at Okayama University Hospital and eight SPs. The trainee dentists conducted initial interviews with the SPs after completing the Japanese version of the Jefferson Scale of Empathy. Their interviews were recorded and analyzed using the Roter Interaction Analysis System. The SPs assessed the trainees’ communication immediately after each interview. The trainee dentists were classified into two groups (more positive and less positive groups) according to SP assessment scores.

**Results:** Compared with the less positive trainees, the more positive trainees scored higher on the [Emotional expression] and lower on the [Medical data gathering] Roter Interaction Analysis System categories. There was no difference in [Dental data gathering] between the two groups. The SPs of more positive trainees had higher rates of [Positive talk] and [Emotional expression] and lower rates of [Medical information giving] and [Dental information giving]. The trainees with more positive ratings from SPs had significantly higher Jefferson Scale of Empathy total scores.

**Conclusion:** The results of this study suggest that responding to the SPs’ emotions is a relevant characteristic of dentist–SP communication to SPs’ positive assessment in medical interviews. Further, trainees’ self-reported empathy was related with the SPs’ assessment of trainees’ communication, which indicated that patient satisfaction can be improved by increasing the dentist’s empathy. Thus, an empathic attitude among dentists is a significant determinant of patient satisfaction.

## Introduction

Effective communication is a critical determinant of delivering better care to patients. Extensive medical literature has suggested that a good relationship between physicians and patients has positive effects on patient outcomes, such as increasing patient satisfaction [1,2], reducing anxiety/distress [3], and increased adherence to treatment [4]. Empathy is also considered to be an important determinant of clinical outcomes. Patients with diabetes who have highly empathic physicians have good metabolic control [5], and patients with HIV who have highly empathic clinicians have higher medication self-efficacy [6].

There is not an exception to this rule in the dental context. Imanaka et al. [7] found in their survey that communication with dentists was the most important determinant of patient satisfaction in a dental hospital. Armfield et al. [8] also identified that the interpersonal characteristics of dentists and staff members, such as friendliness and respectfulness to patients, were the most common influencers of patient satisfaction. Dentists’ negative attitudes in their communication with patients were a significant predictive factor of dental fear [9]. In addition, a negative interpersonal relationship between dentists and patients, such as patients’ negative beliefs and dislike of dentists, was one of the major explanatory factors for missed and cancelled dental appointments [10]. Muirhead et al. [11] found that older people who expressed a lack of trust and confidence in their dentists tended to report worse dental conditions, such as difficulty pronouncing words, painful mouth aches, and feeling uncomfortable while eating. Furthermore, empathy in dentistry also seems to be associated with patient outcomes, especially increasing adherence to and success of treatment. Bernson et al. [12] analyzed interviews with patients who experienced dental-related fear and concluded that empathy was among the main attributes that made dental care accessible to them. Sarnat et al. [13] also found that an empathic approach by dentists was correlated with patients’ cooperative behavior during treatment and the success of treatment.

Good dentist–patient communication is essential to the delivery of dental care. However, research investigating which observable behaviors impact outcomes is scarcely conducted. Further, many studies exploring empathy and patient outcomes in medical settings have been conducted, but fewer have been conducted in dental settings. The aims of this study are to clarify the characteristics of trainee dentists enrolled in a 1-year compulsory postgraduate dental training program and to observe their simulated patient (SP) communication to examine how the level of the trainee dentists’ self-reported empathy influences SPs’ assessment of the trainees’ communication during initial medical interviews.

## Materials and methods

### Participants

A total of 100 trainee dentists (47 male and 53 female) enrolled in a 1-year postgraduate clinical training course at Okayama University Hospital in 2015 and 2016 and eight SPs from the Okayama Working Group for Simulated Patients (one male and seven female) participated. Dental education in Japan consists of a six-year undergraduate program. After acquiring a license, a 1-year postgraduate clinical training program is compulsory.

All trainees provided informed consent in writing after receiving an explanation of this study. Their participation was voluntary and did not influence their evaluation in the program. The Ethics Committee of Okayama University Graduate School of Medicine, Dentistry and Pharmaceutical Sciences approved this study (No. 2219).

### Data collection procedure

The trainee dentists conducted initial interviews with the SPs 3 months after the start of clinical training. The SPs primarily presented concerns about the potential severity of persistent stomatitis on their tongues. The trainee dentists completed the Japanese version of the Jefferson Scale of Empathy (JSE, HP version) immediately before the interviews, which were videotaped and had no time constraints. Immediately after each interview, the SPs assessed the trainees’ communication using an assessment sheet. The medical interviews were implemented once yearly in 2015 and 2016.

The trainee dentists were classified into two groups (i.e., the more positive and less positive groups) based on the median SP assessment score (11.0) of trainees’ communication.

## Measures

### Assessment sheet: SP assessment of trainee dentists’ communication

The assessment sheet (Table 1) consists of five items measured on a 4-point scale (0 = disagree, 1 = disagree somewhat, 2 = agree somewhat, 3 = agree). Total possible scores range from 0 to 15, where a higher score indicates more positive assessment. This assessment sheet was made in reference to the American Board of Internal Medicine’s Patient Assessment survey questionnaire [14].

**Table 1.** Assessment sheet by SPs.

|  | **How was the dental trainee's performance at:**<br>**(circle one number for each item)** | **Agree** | **Agree**<br>**somewhat** | **Disagree**<br>**somewhat** | **Disagree** |
| --- | --- | --- | --- | --- | --- |
| 1 | Listening carefully while you are talking? | 3 | 2 | 1 | 0 |
| 2 | Understanding your worry and uneasiness? | 3 | 2 | 1 | 0 |
| 3 | Speaking with appropriate words and speed in plain language? | 3 | 2 | 1 | 0 |
| 4 | Treating you like you are on the same level; never "talking down" to you or treating you like a child? | 3 | 2 | 1 | 0 |
| 5 | Overall, do you want to see this dental trainee? | 3 | 2 | 1 | 0 |

### TheRoterinteractionanalysissystem(RIAS):Communication characteristics of trainee dentists during medical interviews with SPs

We analyzed the videotaped medical interviews using the Roter interaction analysis system (RIAS), which is a valid and reliable instrument developed to analyze physician–patient interactions during consultations and is currently one of the most widely used systems of its kind in Western countries [18]. The validity and reliability of RIAS has also been examined in the Japanese population [19].

In RIAS, the dialogue between medical professional and patient is divided into “utterances,” defined as the minimum unit comprising one thought or one piece of information. In the Japanese version of RIAS, each utterance is classified into only one of 42 mutually exclusive code categories. In this study, we added 6 new categories to distinguish dental conversations from other medical conversations and then concentrated all categories into 14 larger clusters based on similarity of content (Table 2).

**Table 2.** RIAS categories in this study.

| Cluster | Each category |
| --- | --- |
| Relationship building | Personal remarks, Social conversation, Remediation, Partnership statements, Self-disclosure statement |
| Positive talk | Laughing, Telling jokes, Showing approval-direct, Giving compliments-general, Showing agreement or understanding, Back-channel responses |
| Negative talk | Showing disapproval-direct, Criticizing-general |
| Emotional expression | Empathizing statements, Legitimizing statements, Showing concern or worry, Reassurance, Encouragement or showing optimism, Asking for reassurance |
| Facilitative behaviors | Giving orientation, Instructing, Paraphrasing/checking for understanding, Asking for understanding, Requesting repetition, Asking for opinions, Asking for permission, Transition words, Requesting services or medication |
| Counseling/direction | Counseling or direction about any topic |
| Medical data gathering | Open or Closed questions regarding medical conditions or therapeutic regimen |
| Psychosocial data gathering | Open or Closed questions regarding psychosocial or lifestyle issues |
| Dental data gathering | Open or Closed questions regarding current dental history <sup>a</sup> or past dental history <sup>a</sup> |
| Data gathering about other issues | Open or Closed questions about other issues |
| Medical information giving | Information giving about medical conditions or therapeutic regimen |
| Psychosocial information giving | Information giving about psychosocial or lifestyle issues |
| Dental information giving | Information giving about current dental history<sup>a</sup> or past dental history<sup>a</sup> |
| Information giving about other issues | Information giving about other issues |

<sup>a</sup> New category

RIAS coding is done directly from audio or videotapes rather than transcripts. Therefore, utterances can be categorized based on voice tone and phrasing cues, not only literal meaning.

The main coder (S.W.) analyzed all videotapes according to the RIAS manual [18], and 20% of all of the videotapes (20 tapes, randomly selected) were independently double coded by the second coder (T.Y.) to assess inter-coder reliability. Both coders completed the RIAS coding training offered by RIAS Japan. We calculated the Spearman correlation coefficients between the two coders’ results for all categories with a mean frequency greater than two per medical interview. The average correlation was 0.77 (0.56–0.95) for the trainee dentists and 0.76 (0.61–0.90) for the SPs, which confirmed the reliability of the coding.

The percentage rates of the trainees’ and SPs’ utterances for each category were calculated. The trainees’ and SPs’ overall numbers of utterances were the denominator, and the number of utterances in each category was the numerator, similar to the calculation methods of other studies [20-24]. The percentage rate was used instead of the absolute number of utterances to control for interview length.

### The Jefferson Scale of Empathy (JSE, Health Professional version): Self-evaluation of trainees’ empathy

The Jefferson Scale of Empathy (JSE, Health Professional version) was developed to measure empathy specifically in physicians and health professionals [15]. The reliability and validity of the Japanese version of the JSE has been confirmed [16,17].

The JSE includes 20 items answered on a 7-point Likert-type scale (1 = strongly disagree, 7 = strongly agree) with a total score range of 20–140. Half of the items are reverse scored. A higher score shows a more empathic orientation toward patient care (Table 3).

**Table 3.**
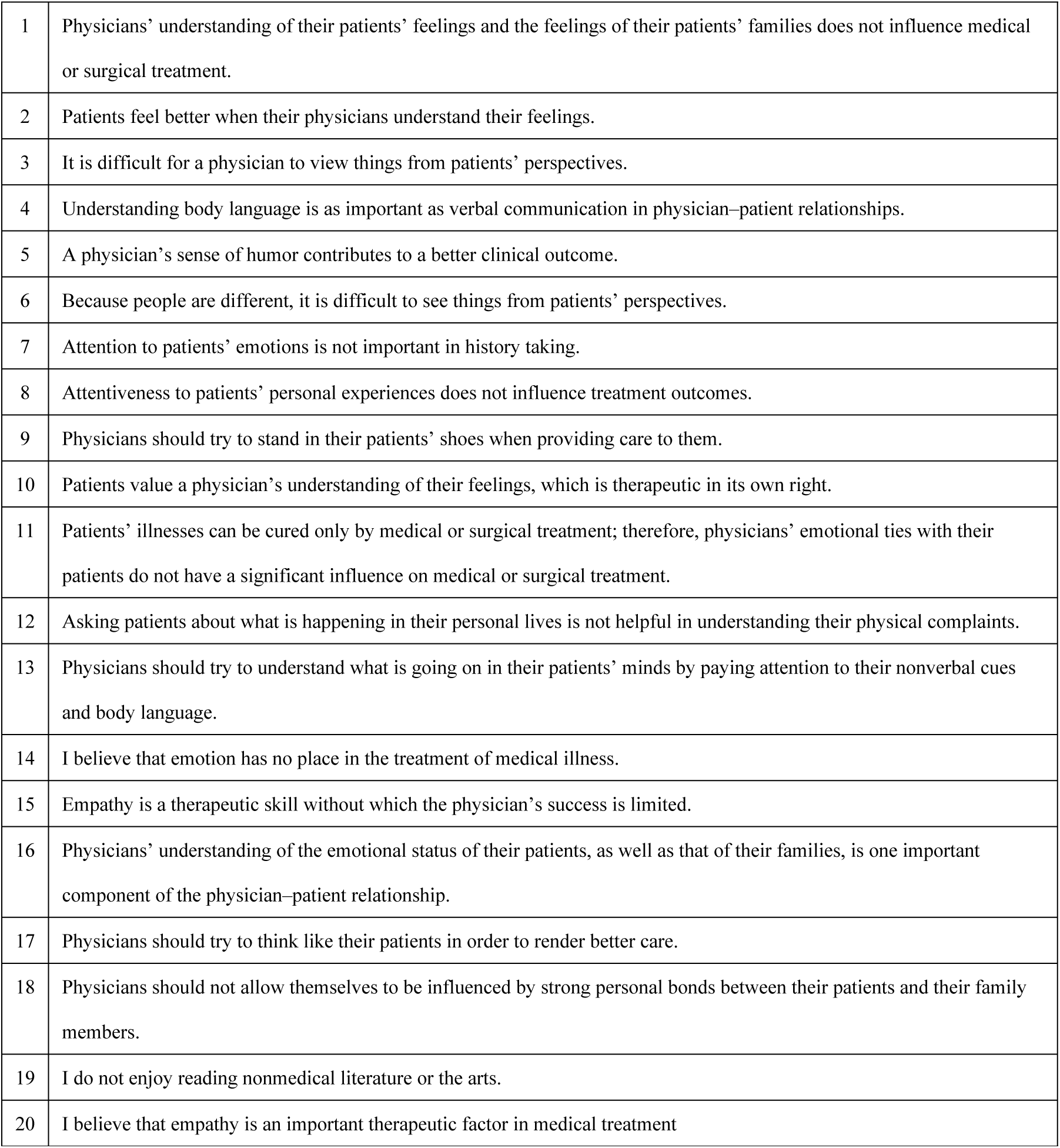
Jefferson Scale of Empathy (JSE, Health Professional version).

|  |  |
| --- | --- |
| 1 | Physicians' understanding of their patients' feelings and the feelings of their patients' families does not influence medical or surgical treatment. |
| 2 | Patients feel better when their physicians understand their feelings. |
| 3 | It is difficult for a physician to view things from patients' perspectives. |
| 4 | Understanding body language is as important as verbal communication in physician–patient relationships. |
| 5 | A physician's sense of humor contributes to a better clinical outcome. |
| 6 | Because people are different, it is difficult to see things from patients' perspectives. |
| 7 | Attention to patients' emotions is not important in history taking. |
| 8 | Attentiveness to patients' personal experiences does not influence treatment outcomes. |
| 9 | Physicians should try to stand in their patients' shoes when providing care to them. |
| 10 | Patients value a physician's understanding of their feelings, which is therapeutic in its own right. |
| 11 | Patients' illnesses can be cured only by medical or surgical treatment; therefore, physicians' emotional ties with their patients do not have a significant influence on medical or surgical treatment. |
| 12 | Asking patients about what is happening in their personal lives is not helpful in understanding their physical complaints. |
| 13 | Physicians should try to understand what is going on in their patients' minds by paying attention to their nonverbal cues and body language. |
| 14 | I believe that emotion has no place in the treatment of medical illness. |
| 15 | Empathy is a therapeutic skill without which the physician's success is limited. |
| 16 | Physicians' understanding of the emotional status of their patients, as well as that of their families, is one important component of the physician–patient relationship. |
| 17 | Physicians should try to think like their patients in order to render better care. |
| 18 | Physicians should not allow themselves to be influenced by strong personal bonds between their patients and their family members. |
| 19 | I do not enjoy reading nonmedical literature or the arts. |
| 20 | I believe that empathy is an important therapeutic factor in medical treatment |

### Statistical analyses

The mean percentages of trainees’ and SPs’ utterances for each category, the total JSE score, and the length of the medical interview were compared between trainee dentists with more and less positive SP assessments. We used the Mann-Whitney test to examine the percentage rates of utterances and unpaired *t*-tests for total JSE scores and the length of the medical interview. All analyses were performed in SPSS (version 23 for Windows, IBM, Tokyo, Japan), and the significance level was 0.05.

## Results

### SP assessment of trainee dentists’ communication

The mean total score was 11.2 (SD = 2.9, range 2–15). The trainee dentists whose SP assessment scores were ≥12 were classified as the more positive group (n = 47), and those who scored <12 were classified as the less positive group (n = 53).

## RIAS

### Percentage rates of the trainees’ and SPs’ utterances

The mean percentage rates of the trainees’ and SPs’ utterances for the clusters are given in Tables 4 and 5, respectively. We express the cluster names in brackets in this study. Compared with the trainee dentists whose SP assessment was less positive, those with more positive assessments had more [Emotional expression], especially empathic and legitimating statements. However, there was a lower proportion of [Medical data gathering] among the more positive trainees. There was no difference in [Dental data gathering] between the two groups.

**Table 4.** Proportions and frequencies of clusters of RIAS categories for trainee dentists with more positive and negative SP assessments.

|  | More positive n = 47 |  | Less positive n = 53 |  | P-value |
| --- | --- | --- | --- | --- | --- |
|  | Median (N) | SD | Median (N) | SD |  |
| Total utterances | 54.95% (115.7) | 4.32% | 55.63% (90.5) | 3.74% | 0.419 |
| Relationship building | 3.75% (4.1) | 1.35% | 4.37% (3.8) | 1.63% | 0.076 |
| Positive talk | 43.15% (50.1) | 7.47% | 41.17% (38.0) | 9.01% | 0.373 |
| Negative talk | 0.00% (0.0) | 0.00% | 0.02% (0.0) | 0.15% | 0.346 |
| Emotional expression | 2.13% (2.6) | 1.90% | 0.93% (0.9) | 1.29% | 0.000** |
| Facilitative behaviors | 26.78% (30.8) | 5.07% | 26.57% (23.9) | 7.03% | 0.841 |
| Counseling/direction | 0.00% (0.0) | 0.00% | 0.00% (0.0) | 0.00% | 1.000 |
| Medical data gathering | 7.30% (8.4) | 2.72% | 8.77% (7.8) | 4.16% | 0.026* |
| Psychosocial data gathering | 1.45% (1.8) | 1.33% | 1.58% (1.4) | 1.36% | 0.584 |
| Dental data gathering | 15.08% (17.5) | 4.48% | 16.42% (14.5) | 4.32% | 0.107 |
| Data gathering about other issues | 0.00% (0.0) | 0.00% | 0.00% (0.0) | 0.00% | 1.000 |
| Medical information giving | 0.15% (0.2) | 0.40% | 0.08% (0.1) | 0.36% | 0.150 |
| Psychosocial information giving | 0.02% (0.0) | 0.12% | 0.01% (0.0) | 0.10% | 0.921 |
| Dental information giving | 0.20% (0.2) | 0.69% | 0.07% (0.1) | 0.26% | 0.554 |
| Information giving about other issues | 0.00% (0.0) | 0.00% | 0.00% (0.0) | 0.00% | 1.000 |

\*P-value <0.05, \*\*P-value <0.01

**Table 5.**
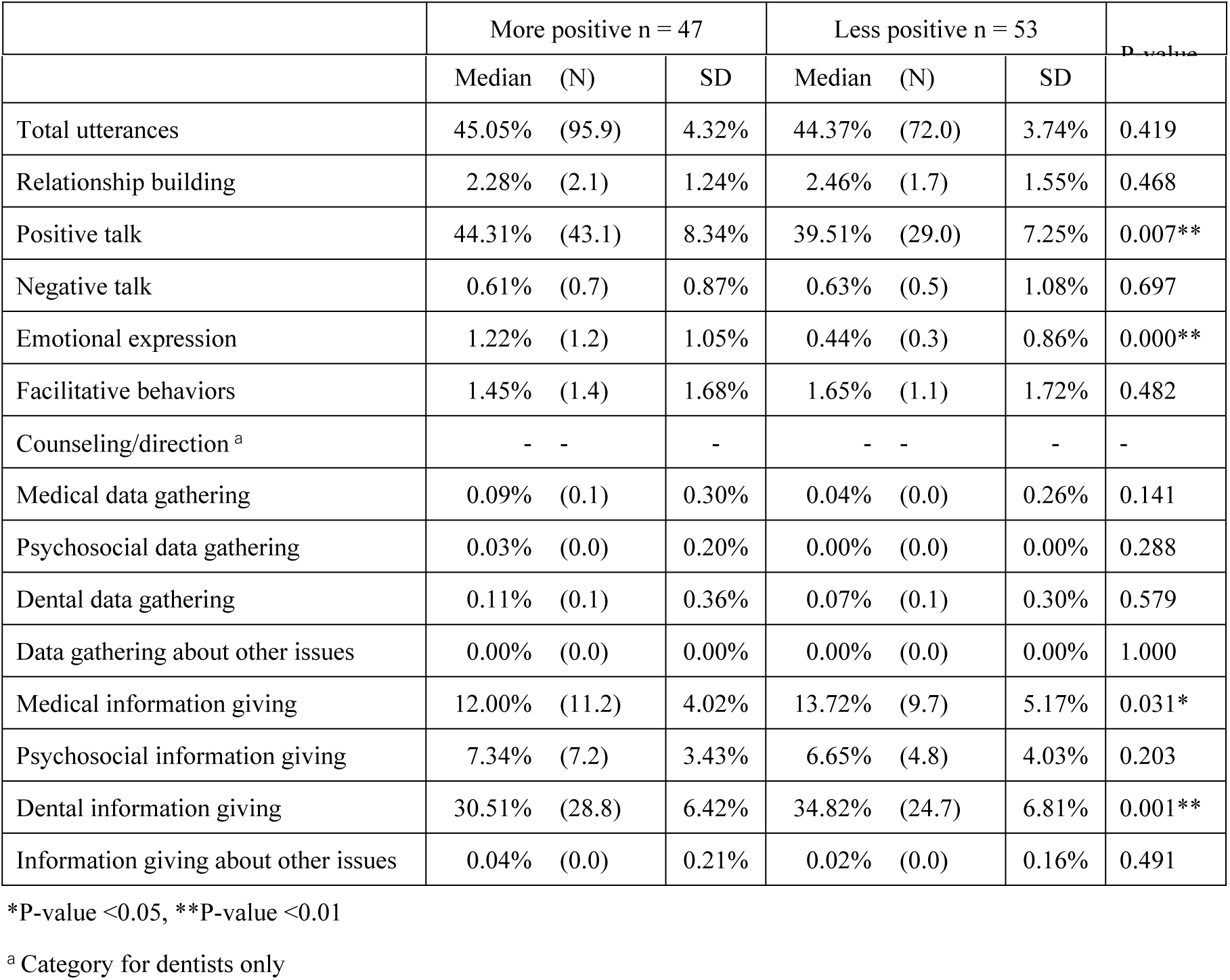
Proportion and frequency of clusters of RIAS categories for SPs who assessed trainee dentists positively and negatively.

Among SPs, [Positive talk], especially back-channel response, and [Emotional expression], which includes concerns, had higher rates in the interviews with more positive trainees. However, the rates of [Medical information giving] and [Dental information giving] were lower in the positive group.

### JSE

The mean total JSE score was 102.00 (64–132, SD = 12.45). There was a significant difference in JSE total scores between the more positive and less positive groups (104.8 vs. 99.5).

### Length of the medical interview

The mean interview length was 8m34s (SD = 2m30s, range 3m43s–17m56s). The mean length of the interviews of more positive trainee dentists was significantly longer than that of the less positive ones (9m34s vs 7m40s).

The results indicate that SPs regarded trainees whose interviews had longer duration more favorably.

## Discussion

In this study, we explored the characteristics of dentist–SP communication that influence SP assessment in initial medical interviews. We also examined how the trainee dentists’ level of self-reported empathy influences their assessments by SPs.

Trainee dentists who received higher ratings from SPs were more emotionally expressive, showing their empathy and legitimizing the SPs’ statements. Their SPs, in turn, expressed their emotions and engaged in positive talk more, revealing that the trainees with higher assessments were able to draw out SPs’ concerns or worries. This suggests that recognizing the SPs’ emotions and expressing acceptance of them with verbal or nonverbal communication leads the SPs to open their minds and be more active conversationally. Such empathizing was received positively by SPs and led to higher assessments of the trainees. This is a reciprocal interaction. Some prior studies have had similar findings to this result. Roter et al. [25] investigated the effects of communication skill training and found that trained doctors used more facilitation and tended to engage in emotional talk more. Their patients also tended to use more positive statements and report higher satisfaction than the patients of untrained doctors did. Dulmen et al. [26] found that more anxious patients preferred empathic doctors and that empathic responses were rated as adequate responses by doctors. The fact that the SPs in our study expressed concerns might reflect this finding.

In the dental context, the relationship between dentist communication and patient satisfaction does not seem to be simple. Sondell et al. [27] reported that dentist communication was associated with patient satisfaction only immediately after a specific visit but not with overall patient satisfaction in the longer term in prosthodontic patients. Even though the specific dimensions of dentist–patient communication that impacted patient satisfaction were not detected in this study, this may imply that observable dentist–patient communication may not be directly related to treatment outcomes, which reflect many characteristics of dentistry. A consultation in dentistry almost always corresponds with a subsequent invasive procedure, and manual skills also affect treatment success, such as in prosthodontics, endodontics, and oral surgery. Patient satisfaction regarding clinical treatment quality has received attention and is often assessed in the dental literature [28]. The SPs in our study assessed dentists’ communication only during their interviews, which may affect our result that dentists with empathic communication received higher assessments from SPs.

With regard to information gathering, the rate of [Medical data gathering] (i.e., questions regarding medical conditions or therapeutic regimen) in trainees with more positive assessments from SPs was significantly lower than that in those who had less positive assessments. Further, there was no difference in the rate of [Dental data gathering] (i.e., questions regarding current and past dental histories) between the two trainee groups. SPs correspondingly shared a lower rate of information about these topics with the more positive trainees. This indicates that trainees with positive assessments were less engaged in [Medical data gathering] and [Dental data gathering]; however, in terms of the number of utterances, the more positive trainees and SPs had more utterances regarding [Medical data gathering] and [Dental data gathering] compared with those in the less positive group, even though this was not statistically tested. Thus, this result does not mean that more positive trainees and their SPs gathered or gave less biomedical information but suggests that they allotted their utterances more to [Emotional expression], such as responding and showing emotions, than gathering and giving information compared with those in the less positive group. Gathering relevant information is indispensable for accurate diagnosis, and responding to patients’ emotions is a relevant characteristic for patient satisfaction. Both aspects are needed to make medical interviews successful.

The three functions of medical interviews are 1) relationship building between health provider and patient, 2) assessment of patients’ problems, and 3) management of patients’ problems [29]. Each function is equally important, but establishing rapport tends to precede data gathering. In the context of dentistry, where verbal communication by patients is often restricted during oral treatment, consultation prior to the treatment plays an important role in the dentist’s understanding of the patient’s psychosocial problems, such as concern, anxiety, and expectation, especially when an invasive procedure is expected. The mouth is a very sensitive part of the body [30], and dental fear/anxiety is widely prevalent and can result in avoidance of dental care [31]. Thus, it is necessary to consider patients’ emotions and to understand their problems, especially during their first visit, when they tend to be nervous.

Both medical and dental information gathering and empathic communication are needed, and therefore, a reasonable amount of time for the interview is required to fulfill both aspects. We found that the interviews of trainee dentists who received more positive assessments from SPs took more time than those of less positive trainees. Although some previous studies showed no association between interview length and patient satisfaction [32,33], some other studies reported that female gender was among the factors related to longer consultation length [34,35]. The fact that the SPs in our study were mostly female might affect our findings.

Our notable findings are that the trainees with higher self-reported empathy received more positive ratings from SPs and that the trainees who received more positive assessments used more empathic communication, indicating that cognitive measures of empathy may reflect behavioral measures. The trainees in this study may have accurately assessed their capability to express their empathy during interviews. Some earlier studies agreed with our findings [36,37]; however, some studies had contradictive results [38,39], which suggests that further studies are needed to confirm our results.

Considering our findings, patient satisfaction can be improved by increasing the dentist’s empathy. As existing literature has indicated effectiveness at enhancing clinicians’ empathy [40], it seems that empathy can be taught and learned. A notable preceding study demonstrated that first year dental students’ humanistic attitude, including that shown in training on patient-centered medical interviews, was associated with 3^rd^-year clinical performance, including patient management and manual skills [41]. Therefore, it is worth fostering an empathic attitude toward patients from the early stages of the dental curriculum. In the case of trainees, as some studies have reported declining empathy during residency [42,43], it is also important to provide feedback on their empathy in patient encounters and provide them with opportunities to review it.

Because our study employed only one institution and had a small sample size, we cannot eliminate the impact of gender concordance between dentists and SPs, which may limit the ability to generalize these findings to a wider population. It would be better to verify these findings in future studies.

## Conclusion

This study provided evidence that responding to patients’ emotions is a relevant characteristic of dentist–patient communication for SP’s positive assessment. For this reason, the conversation during medical interviews in dental settings should not be restricted to biomedical topics but also include responding to patients’ emotional statements. Additionally, trainees’ self-reported cognitive empathy was related with SPs’ assessment of the trainees’ communication. These findings add to a body of literature that indicates that promoting empathic attitudes is a significant aspect of the dental education curriculum.

## Acknowledgments

The authors express their appreciation to all of the trainee dentists and SPs who agreed to participate in the study. The authors also thank the residents who assisted with implementation of the simulated medical interviews. We thank Richard Lipkin, PhD, from Edanz Group (www.edanzediting.com/ac) for editing a draft of this manuscript.

